# Neuro-Environmental Interactions: a time sensitive matter

**DOI:** 10.1101/2023.05.04.539456

**Authors:** Azzurra Invernizzi, Stefano Renzetti, Elza Rechtman, Claudia Ambrosi, Lorella Mascaro, Daniele Corbo, Roberto Gasparotti, Cheuk Y. Tang, Donald R. Smith, Roberto G. Lucchini, Robert O. Wright, Donatella Placidi, Megan K. Horton, Paul Curtin

**Affiliations:** Department of Environmental Medicine and Public Health, Icahn School of Medicine at Mount Sinai, New York, NY, USA; Linus Biotechnology Inc., New York, NY, USA; Department of Medical and Surgical Specialties, Radiological Sciences and Public Health, University of Brescia, Brescia, Italy; Department of Neuroscience,Neuroradiology Unit, ASST Cremona, Italy; ASST Spedali Civili Hospital, Brescia, Italy; Department of Medical Surgical Specialties, Radiological Sciences and Public Health, University of Brescia, Italy; Department of Microbiology and Environmental Toxicology, University of California Santa Cruz, Santa Cruz, CA, USA; Department of Environmental Health Sciences, Robert Stempel School of Public Health, Florida International University, Miami, Florida, USA

## Abstract

The assessment of resting state (rs) neurophysiological dynamics relies on the control of sensory, perceptual, and behavioral environments to minimize variability and rule-out confounding sources of activation during testing conditions. Here, we investigated how temporally-distal environmental inputs, specifically metal exposures experienced up to several months prior to scanning, affect functional dynamics measured using rs functional magnetic resonance imaging (rs-fMRI). We implemented an interpretable XGBoost-Shapley Additive Explanation (SHAP) model that integrated information from multiple exposure biomarkers to predict rs dynamics in typically developing adolescents. In 124 participants (53% females, ages: 13-25 years) enrolled in the Public Health Impact of Metals Exposure (PHIME) study, we measured concentrations of six metals (manganese, lead, chromium, cupper, nickel and zinc) in biological matrices (saliva, hair, fingernails, toenails, blood and urine) and acquired rs-fMRI scans. Using graph theory metrics, we computed global efficiency (GE) in 111 brain areas (Harvard Oxford Atlas). We used a predictive model based on ensemble gradient boosting to predict GE from metal biomarkers, adjusting for age and biological sex. Model performance was evaluated by comparing predicted versus measured GE. SHAP scores were used to evaluate feature importance. Measured versus predicted rs dynamics from our model utilizing chemical exposures as inputs were significantly correlated (*p*< 0.001, *r* = 0.36). Lead, chromium, and copper contributed most to the prediction of GE metrics. Our results indicate that a significant component of rs dynamics, comprising approximately 13% of observed variability in GE, is driven by recent metal exposures. These findings emphasize the need to estimate and control for the influence of past and current chemical exposures in the assessment and analysis of rs functional connectivity.

## Introduction

Intrinsic functional connectivity of the brain has been widely investigated through the analysis of spontaneous (task- independent) blood-oxygen level dependent (BOLD) fluctuations at rest. Resting-state fMRI (rs) allowed the discovery of multiple functional networks underlying cognitive, behavioral and perceptual processing^1,2^ and facilitated further understanding of the temporal and spatial correlation patterns of interconnected brain regions. These correlation patterns are observed in controlled conditions to ensure that variability in sensory, perceptual, or behavioral-evoked neural processing is minimal, thus providing a baseline to characterize connectivity patterns at rest. However, this use of task-free data inherently presumes that the impact of the environment on rs signal is essentially concurrent, or minimally lagged, during data acquisition. Here, we investigated how temporally-distal environmental inputs, specifically metal exposures experienced up to several months prior to testing, affect rs functional dynamics.

To fully characterize the impact of metals exposure on the brain, recent studies have begun investigating the combined or synergistic effect of multiple co-exposures, which may better capture the complex exposure landscape encountered in “real-world” circumstance^3–56^. A challenge in this approach, and in any assessment of exposure, is that differential toxicokinetics involved in varying metal exposures yield a heterogenous distribution of metal biomarkers among different biological media. Therefore, each medium (i.e. blood, urine, etc) provides complementary information on different biological processes. To accommodate this reality, recent studies leverage mixtures-based methods to combine information from multiple exposure matrices^78–10^. These metals mixture-based models demonstrated a more-negative impact on neurodevelopment than single metal model^11–15^. However, these mixture-based statistical approaches require a large sample size to allow appropriate cross-validation procedures, limiting their utility in smaller studies.

In this study, we introduce a novel, alternative and interpretable approach to link high-dimensional environmental exposure assessment to functional connectivity utilizing a machine learning (ML) based predictive framework that allows us to: (a) evaluate the model performance based on the predictive efficacy of the model and (b) leverage all available data simultaneously, much as a mixture approach aims to achieve, while implementing a robust leave-one-out cross validation paradigm to ensure generalizability. The XGBoost model used here has the ability to capture complex interactions among features as well as non-linear relationships between features and classification labels. To help interpretability, a game-theory based measure of variable importance, the Shapley Additive Explanation (SHAP)^16^ method was used to evaluate the importance of each feature in our final model. SHAP scores previously have been applied to explore gene-gene and gene-environment interactions. Here for the first time, we applied SHAP scores to investigate brain-environment dependencies.

We leveraged this platform to explore the importance of temporally-distal environmental chemical inputs on rs intrinsic functional connectivity. Utilizing multi-modal exposure assessment combined with a ML-based modeling platform, we generated a predictive model to determine the extent to which contemporaneous rs functional connectivity could be predicted from environmental inputs experienced up to months prior. Critically, our results suggest that the role of past chemical exposures is a critical variable for future rs studies to control for in the evaluation of rs functional connectivity. These findings highlight the utility of leveraging interpretable machine learning algorithms in neuroexposomic investigations for discovering overlooked interactions between the brain and environmental exposures.

## Results

The complete pipeline of rs fMRI data analysis, model implementation and feature analysis is presented in Figure 1; additional details of the ML model are provided in *SI Appendix*. Observed GE was significantly correlated with predicted GE from the XGBoost model trained with all metal concentrations (Mn, Pb, Cr, Cu, Ni and Zn) measured in blood, urine, hair, saliva, fingernails and toenails as inputs (*p*< 0.001) with a correlation coefficient of 0.36. Based on this correlation, the explained variance of the final model was computed (VE= 13%). To interpret the influence of each feature in the model (i.e., individual metal exposure in each medium) on predicting GE, SHAP scores were calculated to measure feature importance, both at the level of the absolute mean SHAP score (Figure 1b), and in consideration of SHAP scores relative to feature distributions (Figure 1c). These results indicate that urinary and nail Pb, blood Cr, and salivary Cu exposures contributed most to the prediction of global efficiency. Further, analysis of SHAP scores relative to the distribution of urinary Pb (UPb) values (Figure 1c) indicates a negative association between UPb and predicted GE; that is, as UPb increased, the predicted value of GE decreased. In contrast, high blood zinc (BZn) values result in positive SHAP scores, indicating a positive association between BZn and predicted GE.

**Figure 1.**
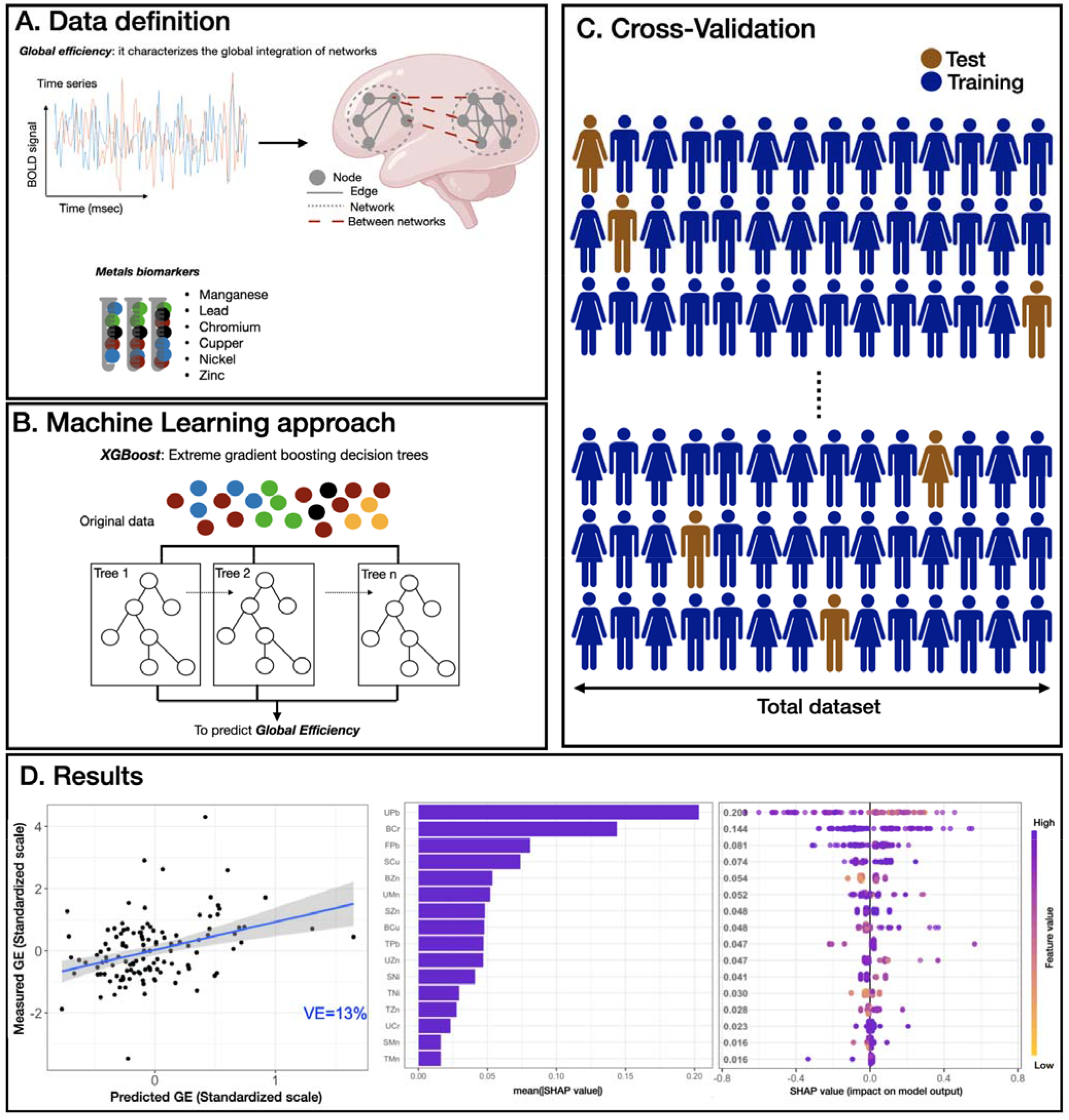
Overview of ML predictive framework. **A**. Resting-state fMRI data were processed and the averaged time-series were extracted using the Harvard-Oxford atlas. Then, the global efficiency (GE) metric was computed for each participant. Small solid gray circles represent nodes of the graphs (brain regions), while gray connecting lines are the edges of the graph (functional connections). Larger dotted circles represent segregated sub-graphs/networks at the whole brain level. For the exposure, six biological samples (blood, saliva, hair, urine, fingernails and toenails) were collected and processed for six metal concentrations (manganese, lead, chromium, copper, nickel and zinc). **B**. XBoost model was used to predict the GE metric using all exposure biomarkers data (“features”). **C**. For model training, 500 iterations of leave-one-out cross validation were used and all features were utilized in the model training. **D**. The performance of the XGBoost model was evaluated by computing the correlation between predicted GE metric versus scaled GE metric (adjusted for age and biological sex) in the hold out subjects obtaining a *p*< 0.001, *r* =0.36 and an explained variance (VE) of 13% (Panel a). Then, SHAP scores were computed for all features (metals exposures) used in the model and the average absolute SHAP value was used to quantify the feature importance. In Panel b, each bar shows the mean absolute SHAP value of each feature, sorted in decreasing order. The most impactful features are displayed and higher SHAP score indicates a more significant contribution in the model prediction, in this case in the prediction of GE metric. Panel c shows the beeswarm plot of SHAP values distribution for the highest ranking features of our model. Feature values associated with a single GE prediction are color-coded, yellow/purple corresponding to low/high metals exposure values, respectively. On the x-axis, the SHAP values are shown representing the impact of a feature with respect to the prediction of the GE metric. Features are sorted using the mean absolute SHAP value in descending order with most important features at the top. Each dot corresponds to one subject. Plots are based on the XBoost model with all features included and leave-one-out-cross validation. BOLD: blood oxygen-level dependent. Features abbreviations: the first letter represents the medium (S=saliva, B=blood, U=urine, H=hair, F=fingernails, T=toenails) and the second and third letters represent the metals (Mn=manganese, Pb=lead, Cr=chromium, Cu=copper, Ni=nickel, Zn=zinc).

## Discussion

We present a novel ML-based framework to evaluate the brain’s distinctive response to temporally-distal environmental inputs, specifically metal exposures occurring up to several months prior to scan. Our findings emphasize that traditional perspectives on environmental control, i.e. homogenization of stimulus and behavioral conditions, fail to account for a critical source of environmentally-driven variance among participants. Leveraging the predictive information provided by the ML model together with SHAP scores, we successfully disentangle, interpret and quantify the strong influence of concurrent and recent past metal exposures that explain 13% of current brain dynamics in adolescents. Finally, this method accurately predicts rs metrics and highlights the power of simultaneously using exposomics data and interpretable ML algorithms for discovering overlooked interactions between environmental exposures and the brain.

Environmental neuroscientists typically assess brain-environmental interactions by investigating the association between individual components of exposure, in our case environmental metals, and brain metrics^17,18^. Based on their unique chemical properties and similar neurobiological mechanisms of actions, several studies report synergistic neurotoxic effects of metals-exposure^19^. Metals within our mixture have been shown to produce such synergistic neurotoxic effects^20–22^, and epidemiological studies suggest that co-exposure to multiple metals, compared to individuals, increases disruption to human neurodevelopment^9,13,14,23^. Few neuroimaging studies account for this synergistic action of multiple metals mixture on neurophysiological dynamics^15^ and rather focus on single-metal assessments^3,4^. Here, we applied a multi-modal exposure assessment (i.e., multiple metals in several media) combined with ML-based modeling to investigate the impact of mixed metal exposures on brain dynamics while retaining the discrete information regarding each individual metal.

Our findings reveal a clear link between temporally-distal environmental exposures and current neurophysiological dynamics. When acquiring rs dynamics to assess brain activation patterns, we typically control for sensory and perceptual environments during testing conditions by removing all external stimuli (room lights or visual stimuli) and encouraging the participant to remain still and relaxed and either keep eyes closed or focused on a non-stimulation fixation point (i.e., seascape or cross). Notably, these co-occurring sensory and perceptual inputs operate on the brain signals within timescales of millisecond-to-second intervals as shown in task-based fMRI data^24^. In contrast, here we show that chemical environmental inputs, in particular exposures that have occurred weeks and months previously, are also important determinants of neurophysiological dynamics. These results may be particularly relevant in the context of the current reproducibility crisis; that is, neuroimaging studies suffer from high variability and lack of reproducibility^25^. This challenge may be explained, at least partially, by addressing the impact of overlooked external stressors (i.e. sociodemographic metrics, -omics data, environmental factors) on the functional brain signals. In this study, we confirm that the past chemical environment is certainly critical to control or account for. Accordingly, future studies should consider how environmental, social and other past exposures might play a role in shaping the recorded brain signals.

Given the high dimensionality of multiple exposures, compounded by multiple exposure media, other typical approaches might be considered (i.e., mixtures-based). More classical mixtures-based approaches (i.e., BKMR^7^, WQS^26–28^) assess multiple mixtures simultaneously and are applied in an explanatory setting, i.e. either to test hypotheses, or to estimate the effects of contributing chemicals. These methods require a large sample to allow appropriate cross-validation procedures and have not been leveraged in a predictive framework to assess the extent to which past metals co-exposures drive the contemporaneous underlying brain dynamics. An additional strength of our approach is the combination of XGBoost and SHAP scores allowing us to explain the contribution of each feature at both a global level, and at a fine-scale relative to the distribution of each measurement. For example, the distribution of SHAP scores for urinary Pb (UPb), the top contributor to rs dynamics prediction, relative to the observed values of UPb indicated a non-linear association, with high UPb values contributing disproportionally to model efficacy. Globally, UPb values were negatively associated with global efficiency, consistent with previous studies showing high lead levels disrupt neuronal activity^15^ and are associated with altered structural connectivity and functional activation patterns in children and adults^29^.

Despite the accuracy and interpretation of the ML-based framework presented here, there are several limitations. Our results show a significant correlation between the observed and predicted rs signal from the XGBoost model that trained with all available metal concentrations. However, in this study, we considered only exposure to six metals. Future studies should also consider examining the effect of additional exposures and omics data (i.e., epigenomics, proteomics, transcriptomics and metabolomics) and sociodemographic metrics to capture the social and physical environment and their impact on the functional brain signals. Given the small sample size of our dataset and the limited exposure assessment, it was not possible to validate with independent and external cohorts to further generalize these results. To account for this, we implemented a leave-one-out cross validation in our model. Finally, this study focuses exclusively on resting state dynamics, these brain-environment interactions may influence task-based functional signals as well.

Utilizing multi-modal exposure assessment combined with a ML-based modeling, we were able to quantify the impact of the temporally-distal environmental on current neurophysiological dynamics. Our work highlights how this continuous brain-environment interaction is key to advance our understanding of neural mechanisms and can inform on both disease pathogenesis and future public health policies.

## Materials and Methods

### Participants

The Public Health Impact of Metal Exposure (PHIME) cohort investigates associations between metal exposure from anthropogenic emissions and developmental health outcomes in adolescents and young adults living proximate to the ferro-manganese industry in northern Italy. Details of the study have been described elsewhere ^30,31^. Inclusion criteria were: birth in the areas of interest; family residence in Brescia for at least two generations; residence in the study areas since birth. Exclusion criteria were: having a severe neurological, hepatic, metabolic, endocrine or psychiatric disorder; using medications (in particular with neuro-psychological side effects); having clinically diagnosed motor deficits or cognitive impairment and having visual deficits that are not adequately corrected. Detailed description of this recruitment process and study design can be found in previous publications ^31,32^. A convenience-based sample of 202 participants (53% female, ages 13-25 years) were selected and willing to participate in a multimodal magnetic resonance imaging (MRI) study, PHIME-MRI. They completed multimodal MRI scans, neuropsychological tests, including measures of IQ (Kaufman Brief Intelligence Test, Second Edition (KBIT-2))^33–35^, memory and motor functions. All participants satisfied eligibility criteria for MRI scanning (i.e., no metal implants or shrapnel, claustrophobia, no prior history of traumatic brain injury, body mass index (BMI) ≤ 40). Manganese, lead, chromium, copper, nickel and zinc (Mn, Pb, Cr, Cu, Ni, Zn, respectively) were measured in saliva, hair, fingernails, toenails, blood and urine, for each PHIME-MRI participant. Complete exposure data (i.e., all metals in all media for a total of 6 components), MRI and covariates data were available for 124 participants (57% females; with an average age of 19.04 years, range=13-25) included in this analysis (69 missing biomarkers and 9 poor MRI quality).

Written informed consent was obtained from parents, while participants provided written assent. Study procedures were approved by the Institutional Review Board of the University of California, Santa Cruz, the ethical committees of the University of Brescia, and the Icahn School of Medicine at Mount Sinai. Details are provided in *SI Appendix*.

### Biomarker measures of exposure

Biological samples including venous whole blood, spot urine, saliva, hair, fingernails and toenails were collected from each subject upon enrollment, as described in detail in previous studies ^36–39^. Biological samples were processed and analyzed for metal concentrations using magnetic sector inductively coupled plasma mass spectroscopy (Thermo Element XR ICP-MS), as described elsewhere ^30,37–39^. A complete overview of biomarkers can be found in Table 1, while Pearson’s correlations between biomarkers is reported in Figure 1. Samples with values less than the LOD were substituted with LOD/square root of 2^40^.

**Table 1.** Metal concentrations (Mn, Pb, Cr, Cu, Ni and Zn) measured in blood, urine, hair, saliva, fingernails and toenails collected from 124 adolescent participants included in the PHIME-MRI study.

| Medium* | Metal | GM | GSD | %>LOD | LOD mean ± SE |
| --- | --- | --- | --- | --- | --- |
| Saliva | Lead | 0.181 | 3.386 | 89 | 0.052 ± 0.004 |
| | Chromium | 0.393 | 3.729 | 90 | 0.125 ± 0.005 |
| | Manganese | 3.178 | 3.084 | 100 | 0.084 ± 0.001 |
| | Nickel | 1.204 | 3.014 | 93 | 0.198 ± 0.011 |
| | Copper | 8.653 | 2.403 | 99 | 0.418 ± 0.034 |
| | Zinc | 45.865 | 2.898 | 100 | 2.3189 ± 0.136 |
| Hair | Lead | 0.108 | 2.744 | 100 | 0.003 ± 0.001 |
| | Chromium | 0.044 | 2.542 | 100 | 0.004 ± 0.001 |
| | Manganese | 0.072 | 2.557 | 100 | 0.006 ± 0.001 |
| | Nickel | 0.071 | 4.024 | 88 | 0.014 ± 0.001 |
| | Copper | 10.190 | 1.629 | 100 | 0.048 ± 0.003 |
| | Zinc | 85.750 | 1.619 | 100 | 0.209 ± 0.008 |
| Fingernails | Lead | 0.030 | 3.326 | 96 | 0.003 ± 0.001 |
| | Chromium | 0.082 | 2.348 | 99 | 0.004 ± 0.001 |
| | Manganese | 0.097 | 2.244 | 99 | 0.006 ± 0.001 |
| | Nickel | 0.192 | 5.522 | 94 | 0.017 ± 0.001 |
| | Copper | 2.813 | 1.523 | 100 | 0.046 ± 0.002 |
| | Zinc | 39.818 | 1.702 | 100 | 0.209 ± 0.009 |
| Toenails | Lead | 0.058 | 2.530 | 100 | 0.003 ± 0.001 |
| | Chromium | 0.113 | 3.113 | 100 | 0.004 ± 0.001 |
| | Manganese | 0.096 | 3.081 | 99 | 0.006 ± 0.001 |
| | Nickel | 0.073 | 3.756 | 80 | 0.017 ± 0.001 |
| | Copper | 2.475 | 1.352 | 100 | 0.046 ± 0.003 |
| | Zinc | 26.665 | 1.763 | 100 | 0.208 ± 0.009 |
| Urine | Lead | 0.338 | 1.980 | 100 | 0.063 ± 0.003 |
| | Chromium | 0.339 | 2.702 | 97 | 0.101 ± 0.004 |
| | Manganese | 0.259 | 3.043 | 85 | 0.115 ± 0.003 |
| | Nickel | 1.193 | 2.566 | 97 | 0.231 ± 0.007 |
| | Copper | 5.604 | 1.800 | 100 | 0.325 ± 0.012 |
| | Zinc | 368.526 | 1.942 | 100 | 4.241 ± 0.302 |
| Blood | Lead | 8.409 | 1.509 | 100 | 0.161 ± 0.004 |
| | Chromium | 0.442 | 4.287 | 75 | 0.205 ± 0.011 |
| | Manganese | 8.364 | 1.509 | 100 | 0.482 ± 0.023 |
| | Nickel | 1.901 | 0.136 | 81 | 0.791 ± 0.035 |
| | Copper | 574.459 | 1.310 | 100 | 1.108 ± 0.037 |
| | Zinc | 3599.253 | 1.448 | 100 | 7.651 ± 0.636 |

**Note:** GM = Geometric mean; GSD = Geometric standard deviation; LOD = limit of detection; SE = standard error of the mean. \*Metrics used to measure metal concentrations within each medium were: blood and saliva (ng/mL), hair (ug/g), urine (ug/mL).

**Figure 1.**
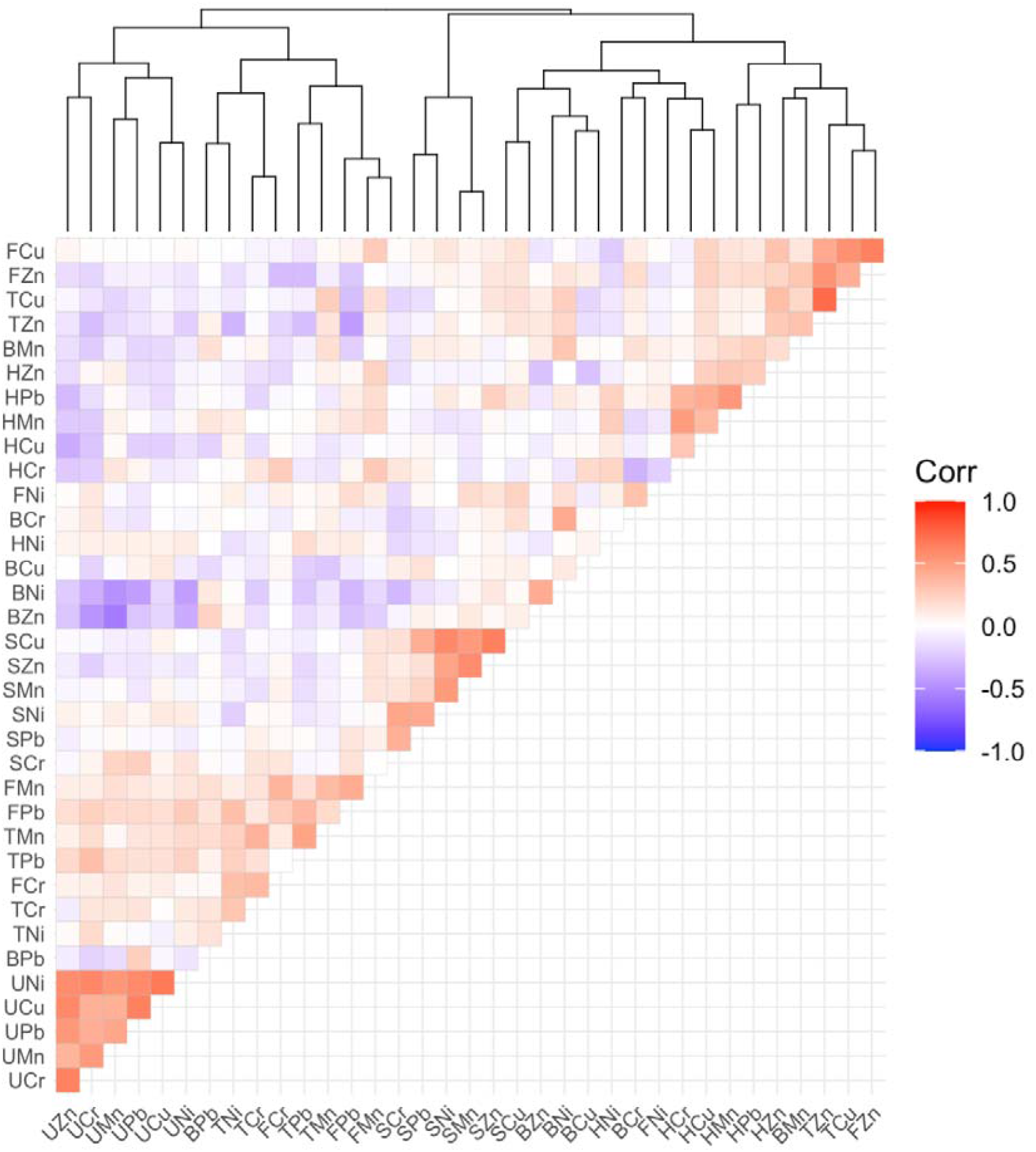
Heatmap of metals exposure. Spearman’s correlations and hierarchical clustering between all biomarkers collected in PHIME-MRI. Components abbreviations: the first letter represents the medium (S=saliva, B=blood, U=urine, H=hair, F=fingernails, T=toenails) and the second and third letters represent the metals (Mn=manganese, Pb=lead, Cr=chromium, Cu=copper, Ni=nickel, Zn=zinc).

### MRI and fMRI data acquisition

Magnetic resonance imaging (MRI) and functional MRI (fMRI) data acquisition was performed on a high-resolution 3-Tesla SIEMENS Skyra scanner using a 64-channel phased array head and neck coil, at the Neuroimaging Division of ASST Spedali Civili Hospital of Brescia. For each participant, a high-resolution 3D T1-weighted structural scan was acquired using a MPRAGE sequence (TR =2400 ms, TE= 2.06 ms, TI=230 ms, acquisition matrix=256×256 and 224 sagittal slices with final voxel size=0.9 mm^3^). Fifty contiguous oblique-axial sections were used to cover the whole brain where the first four images were discarded to allow the magnetization to reach equilibrium. For each subject, a single 10-minute continuous functional sequence using a T2*weighted echo-planar imaging (EPI) sequence (TR=1000 ms, TE=27 ms, 70 axial slices, 2.1 mm thickness, matrix size 108×108, covering the brain from vertex to cerebellum) was acquired. During resting-state scans, lights of the MRI room were off, and participants were instructed to stay awake, relax and daydream (not think about anything) while keeping eyes open. They were presented with an image of a night skyline figure projected on a MRI compatible monitor. Padding was used for comfort and reduction of head motion. Earplugs were used to reduce noise. Data were read by a board-certified radiologist to determine quality and possible incidental findings - no findings were reported.

### fMRI data analyses

Image pre-processing, global efficiency calculation, and statistical analyses were performed using SPM12 (Wellcome Department of Imaging Neuroscience, London, UK), Brain Connectivity toolbox ^41,42^ and customized scripts, implemented in MatLab 2016b (The Mathworks Inc., Natick, Massachusetts) and R (v3.4).

#### Image preprocessing

For each subject, the structural magnetic resonance image was co-registered and normalized against the Montreal Neurological Institute (MNI) template and segmented to obtain white matter (WM), gray matter (GM) and cerebrospinal fluid (CSF) probability maps in the MNI space. FMRI data were spatially realigned, co-registered to the MNI-152 EPI template and subsequently normalized utilizing the segmentation option for EPI images in SPM12. All normalized data were denoised using ICA-AROMA ^43^. Additionally, spatial smoothing was applied (8 millimeters) to the fMRI data. No global signal regression was applied. Based on the Harvard-Oxford ^44^ atlas, 111 regions of interest (ROI; 48 left and 48 right cortical areas; 7 left and 7 right subcortical regions and 1 brainstem) were defined. In this atlas, the brain areas were defined using T1-weighted images of 21 healthy male and 16 healthy female subjects (ages 18-50). The T1-weighted images were segmented and affine-registered to MNI152 space using FLIRT (FSL), and the transformations then applied to the individual brain areas’ label. For each ROI, a time-series was extracted by averaging across voxels per time point. To facilitate statistical inference, data were “pre-whitened” by removing the estimated autocorrelation structure in a two-step generalized linear model (GLM) procedure ^45,46^. In the first step, the raw data were filtered against the 6 motion parameters (3 translations and 3 rotations). Using the resulting residuals, the autocorrelation structures present in the data were estimated using an Auto-Regressive model of order 1 (AR(1)) and then removed from the raw data. Next, the realignment parameters, white matter (WM) and cerebrospinal fluid (CSF) signals were removed as confounders on the whitened data.

#### Network analysis

Global Efficiency (GE) was computed using the Brain Connectivity toolbox ^41,42^ on the defined ROI time course data per subject. GE builds on the concept of efficient integration of communication in a network at a global level. GE is defined as the inverse of the average characteristic path length between all nodes in the networks ^47,48^. For each individual node, with each node defined as an ROI, the shortest number of intermediary nodes required to traverse a path from one node to another was computed. Then, the average number of shortest steps to all defined nodes was computed separately for each node. To correct for the total number of connections between nodes, the inverse of the average number of shortest steps for each node was summed across all network nodes and normalized. GE was scaled from 0 to 1, with a value of 1 indicating maximum GE in observed distribution.

### Statistical analysis

#### Descriptive statistics

Visual inspection and descriptive statistics (geometric mean, geometric standard deviation, and Spearman’s correlation) were used to characterize the metal concentrations in different media.

#### Predictive Modeling

The goal of the predictive modeling was to utilize descriptive statistics (“features”) generated in the descriptive analysis of exposure biomarkers to predict GE metric. The model utilized for predictive classification was a form of ensemble gradient boosting^49^, referred to as XGBoost (“Extreme Gradient Boosting”). This approach was selected for the utility of tree-based models for capturing non-linear dependencies and interactions among features, while also leveraging the efficacy of gradient boosting. For hyperparameter tuning, 500 iterations of leave-one-out cross validation were implemented to evaluate the best-performing set of hyperparameters. Following this, the optimal hyperparameter set was used to train a model with leave-one-out holdout validation, and the performance of this model evaluated by computing the correlation between predicted GE metric versus measured GE metric in each hold out subject. Based on the Pearson correlation coefficient (r), we finally calculated the explained variance of the model (VE).

#### Features importance analysis

Given the ensemble decision trees used in the XGBoost classification algorithm, one feature can be used in multiple locations across the decision trees algorithm making it challenging to interpret the feature importance of the model. For this reason, the Shapley Additive Explanation (SHAP)^16^ method was used to evaluate each feature’s importance in our final model. Based on the trained model, a unique SHAP score was estimated to quantify the contribution of each measurement to model predictions. This score quantifies the effect of each feature on the classification model by measuring the deviation from the average prediction brought by the value of a specific feature. This approach allows the evaluation of non-linear aspects of feature importance; for example, if a given feature contributes to predictive efficacy primarily in cases of extreme scores. In contrast, we subsequently computed the average absolute SHAP value for each feature to capture global importance.

#### Software Implementation

All descriptive statistical analyses and predictive modelling were implemented using R (version 4.2.1) programming language. The following libraries were used: “data.table” and “imputeTS” for data manipulation; “mlr” and “xgboost” for model training, fitting, and prediction; “SHAPforxgboost” and “ggplot2” libraries were used to quantify and visualize model prediction by computing SHAP scores for each feature, respectively.

## Acknowledgements

The authors would like to acknowledge support from the National Institutes of Environmental Health Sciences (NIEHS) grants numbers R01 ES019222, R01 ES013744, P30ES023515, and the European Union through its Sixth Framework Programme for RTD (contract number FOOD-CT-2006-016253).

## Notes

<u>Conflict of Interest:</u> None.

### Competing Interest Statement

The authors have declared no competing interest.

## References

1. Greicius Krasnow, B., Reiss, A. L. & Menon, V. Functional connectivity in the resting brain: a network analysis of the default mode hypothesis. Proc. Natl. Acad. Sci. U. S. A. 100, (2003).

2. Park, H. J. & Friston, K. Structural and functional brain networks: from connections to cognition. Science 342, (2013).

3. de Water, E. et al. Early-life dentine manganese concentrations and intrinsic functional brain connectivity in adolescents: A pilot study. PLoS One 14, (2019).

4. Prenatal manganese exposure and intrinsic functional connectivity of emotional brain areas in children. Neurotoxicology 64, 85–93 (2018).

5. Thomason, M. E. et al. Prenatal lead exposure impacts cross-hemispheric and long-range connectivity in the human fetal brain. Neuroimage 191, (2019).

6. Bauer, J. A. et al. Associations of a Metal Mixture Measured in Multiple Biomarkers with IQ: Evidence from Italian Adolescents Living near Ferroalloy Industry. Environ. Health Perspect. (2020) doi:10.1289/EHP6803.

7. Bobb, J. F. et al. Bayesian kernel machine regression for estimating the health effects of multi-pollutant mixtures. Biostatistics 16, (2015).

8. Critical windows of susceptibility in the association between manganese and neurocognition in Italian adolescents living near ferro-manganese industry. Neurotoxicology 87, 51–61 (2021).

9. Integrated measures of lead and manganese exposure improve estimation of their joint effects on cognition in Italian school-age children. Environ. Int. 146, 106312 (2021).

10. Levin-Schwartz, Y. et al. Multi-media biomarkers: Integrating information to improve lead exposure assessment. Environ. Res. 183, (2020).

11. Horton, M. K. et al. Dentine biomarkers of prenatal and early childhood exposure to manganese, zinc and lead and childhood behavior. Environ. Int. 121, (2018).

12. Freire, C. et al. Prenatal co-exposure to neurotoxic metals and neurodevelopment in preschool children: The Environment and Childhood (INMA) Project. Sci. Total Environ. 621, (2018).

13. Claus, H. B. et al. Associations of early childhood manganese and lead coexposure with neurodevelopment. Environ. Health Perspect. 120, (2012).

14. Claus, H. B., Coull, B. A. & Wright, R. O. Chemical mixtures and children’s health. Curr. Opin. Pediatr. 26, (2014).

15. Invernizzi, A. et al. Topological network properties of resting-state functional connectivity patterns are associated with metal mixture exposure in adolescents. Front. Neurosci. 17, (2023).

16. Failure mode and effects analysis of RC members based on machine-learning-based SHapley Additive exPlanations (SHAP) approach. Eng. Struct. 219, 110927 (2020).

17. Berman, M. G., Kardan, O., Kotabe, H. P., Nusbaum, H. C. & London, S. E. The promise of environmental neuroscience. Nature human behaviour 3, (2019).

18. Berman, M. G., Stier, A. J. & Akcelik, G. N. Environmental neuroscience. Am. Psychol. 74, (2019).

19. de Andrade, L. V., Dos Santos AP M. & Aschner, M. NEUROTOXICITY OF METAL MIXTURES. Advances in neurotoxicology 5, (2021).

20. Tao, S., Liang, T., Cao, J., Dawson, R. W. & Liu, C. Synergistic effect of copper and lead uptake by fish. Ecotoxicol. Environ. Saf. 44, (1999).

21. Lu, C., Svoboda, K. R., Lenz, K. A., Pattison, C. & Ma, H. Toxicity interactions between manganese (Mn) and lead (Pb) or cadmium (Cd) in a model organism the nematode C. elegans. Environ. Sci. Pollut. Res. Int. 25, (2018).

22. Chen, P., Miah, M. R. & Aschner, M. Metals and Neurodegeneration. F1000Res. 5, (2016).

23. Kim, Y. et al. Co-exposure to environmental lead and manganese affects the intelligence of school-aged children. Neurotoxicology 30, (2009).

24. Katwal, S. B., Gore, J. C., Christopher Gatenby, J. & Rogers, B. P. Measuring Relative Timings of Brain Activities Using FMRI. Neuroimage 0, 436 (2013).

25. Marek, S. et al. Reproducible brain-wide association studies require thousands of individuals. Nature 603, 654–660 (2022).

26. Carrico, C., Gennings, C., Wheeler, D. C. & Factor-Litvak, P. Characterization of Weighted Quantile Sum Regression for Highly Correlated Data in a Risk Analysis Setting. J. Agric. Biol. Environ. Stat. 20, 100–120 (2014).

27. Repeated holdout validation for weighted quantile sum regression. MethodsX 6, 2855–2860 (2019).

28. [No title]. https://www.tandfonline.com/doi/full/10.1080/03610918.2019.1577971.

29. Bressler, J. P. & Goldstein, G. W. Mechanisms of lead neurotoxicity. Biochem. Pharmacol. 41, (1991).

30. Lucas, E. L. et al. Impact of ferromanganese alloy plants on household dust manganese levels: implications for childhood exposure. Environ. Res. 138, (2015).

31. Lucchini, R. G. et al. Tremor, olfactory and motor changes in Italian adolescents exposed to historical ferro-manganese emission. Neurotoxicology 33, (2012).

32. Lucchini, R. G. et al. Inverse association of intellectual function with very low blood lead but not with manganese exposure in Italian adolescents. Environ. Res. 118, (2012).

33. Reynolds, C. R., Vannest, K. J. & Fletcher-Janzen, E. Encyclopedia of Special Education, 4 Volume Set: A Reference for the Education of Children, Adolescents, and Adults Disabilities and Other Exceptional Individuals. (Wiley, 2014).

34. Reynolds, C. R., Vannest, K. J. & Fletcher-Janzen, E. Encyclopedia of Special Education, Volume 4: A Reference for the Education of Children, Adolescents, and Adults Disabilities and Other Exceptional Individuals. (John Wiley & Sons, 2018).

35. Kaufman, A. S. & Kaufman, N. L. Kaufman Brief Intelligence Test, Second Edition. Encyclopedia of Special Education Preprint at https://doi.org/10.1002/9781118660584.ese1325 (2014).

36. Lucas, E. L. et al. Impact of ferromanganese alloy plants on household dust manganese levels: implications for childhood exposure. Environ. Res. 138, (2015).

37. Butler, L. et al. Assessing the contributions of metals in environmental media to exposure biomarkers in a region of ferroalloy industry. J. Expo. Sci. Environ. Epidemiol. 29, (2019).

38. Eastman, R. R., Jursa, T. P., Benedetti, C., Lucchini, R. G. & Smith, D. R. Hair as a biomarker of environmental manganese exposure. Environ. Sci. Technol. 47, (2013).

39. Smith, D. et al. Biomarkers of Mn exposure in humans. Am. J. Ind. Med. 50, (2007).

40. Ganser, G. H. & Hewett, P. An accurate substitution method for analyzing censored data. J. Occup. Environ. Hyg. 7, (2010).

41. Rubinov, M. & Sporns, O. Complex network measures of brain connectivity: uses and interpretations. Neuroimage 52, 1059–1069 (2010).

42. Rubinov, M., Kötter, R., Hagmann, P. & Sporns, O. Brain connectivity toolbox: a collection of complex network measurements and brain connectivity datasets. NeuroImage vol. 47 S169 Preprint at https://doi.org/10.1016/s1053-8119(09)71822-1 (2009).

43. Pruim, R. H. R., Mennes, M., Buitelaar, J. K. & Beckmann, C. F. Evaluation of ICA-AROMA and alternative strategies for motion artifact removal in resting state fMRI. Neuroimage 112, 278–287 (2015).

44. Desikan, R. S. et al. An automated labeling system for subdividing the human cerebral cortex on MRI scans into gyral based regions of interest. Neuroimage 31, (2006).

45. Monti, M. M. Statistical Analysis of fMRI Time-Series: A Critical Review of the GLM Approach. Front. Hum. Neurosci. 5, 28 (2011).

46. Bright, M. G. & Murphy, K. Is fMRI ‘noise’ really noise? Resting state nuisance regressors remove variance with network structure. Neuroimage 114, 158–169 (2015).

47. Latora, V. & Marchiori, M. Efficient behavior of small-world networks. Phys. Rev. Lett. 87, (2001).

48. Bullmore, E. & Sporns, O. The economy of brain network organization. Nat. Rev. Neurosci. 13, 336–349 (2012).

49. Chen, T. & Guestrin, C. XGBoost: A Scalable Tree Boosting System. (2016) doi:10.48550/arXiv.1603.02754.

